# The *Cucumis sativus* kinome: Identification, annotation, and expression patterns in response to powdery mildew infection

**DOI:** 10.1101/2023.03.16.532963

**Authors:** Francisco Cleilson Lopes Costa, Welison Andrade Pereira

**Affiliations:** Macaé Ecological Research Center, Institute of Biodiversity and Sustainability, Universidade Federal do Rio de Janeiro (UFRJ), Macaé, RJ, Brazil; Department of Biology, Institute of Natural Sciences, Universidade Federal de Lavras (UFLA), Lavras, MG, Brazil

**Keywords:** Gene expression, Kinase gene family, Metabolism regulation, Plant immunity, Protein evolution, Response to disease

## Abstract

It is widely known that protein kinases (PKs) play a fundamental role in regulating various metabolic processes in plants, from development to response to the environment. However, a detailed characterization of this superfamily is still lacking for several species, such as cucumber (*Cucumis sativus*), especially regarding their involvement in the response to Powdery Mildew (PM) caused by *Podosphaera xanthii*. This study aimed to characterize the cucumber PK family, shedding light on its genomic distribution, classification, and expression patterns triggered by *P. xanthii*. The hidden Markov models (HMMs) analysis uncovered 835 PKs in the cucumber kinome, distributed across its seven chromosomes, and categorized into 20 distinct groups and 123 families, with the RLK group being the most abundant. Evidence of tandem duplication of PK genes was also observed, enriching our understanding of cucumber PKs. To investigate the expression profiles of PK genes in cucumber, we analyzed the transcription levels of all 835 PK genes in RNA-seq data from leaves of resistant and susceptible cultivars of cucumber to *P. xanthii*, which were artificially inoculated. Depending on the treatment, DEGs ranged from 319 to 1,690, with PK DEGs ranging from 8 to 105. The number of PK DEGs varied between the different contrasts analyzed. Notably, we observed a greater number of PK DEGs in susceptible genotypes when challenged by the pathogen. Our findings indicate the role of specific cucumber PKs in regulating metabolic processes in the context of plant-pathogen interactions and pave the way for further research into the intricate mechanisms underlying cucumber responses to Powdery Mildew.

**Graphical Abstract:** 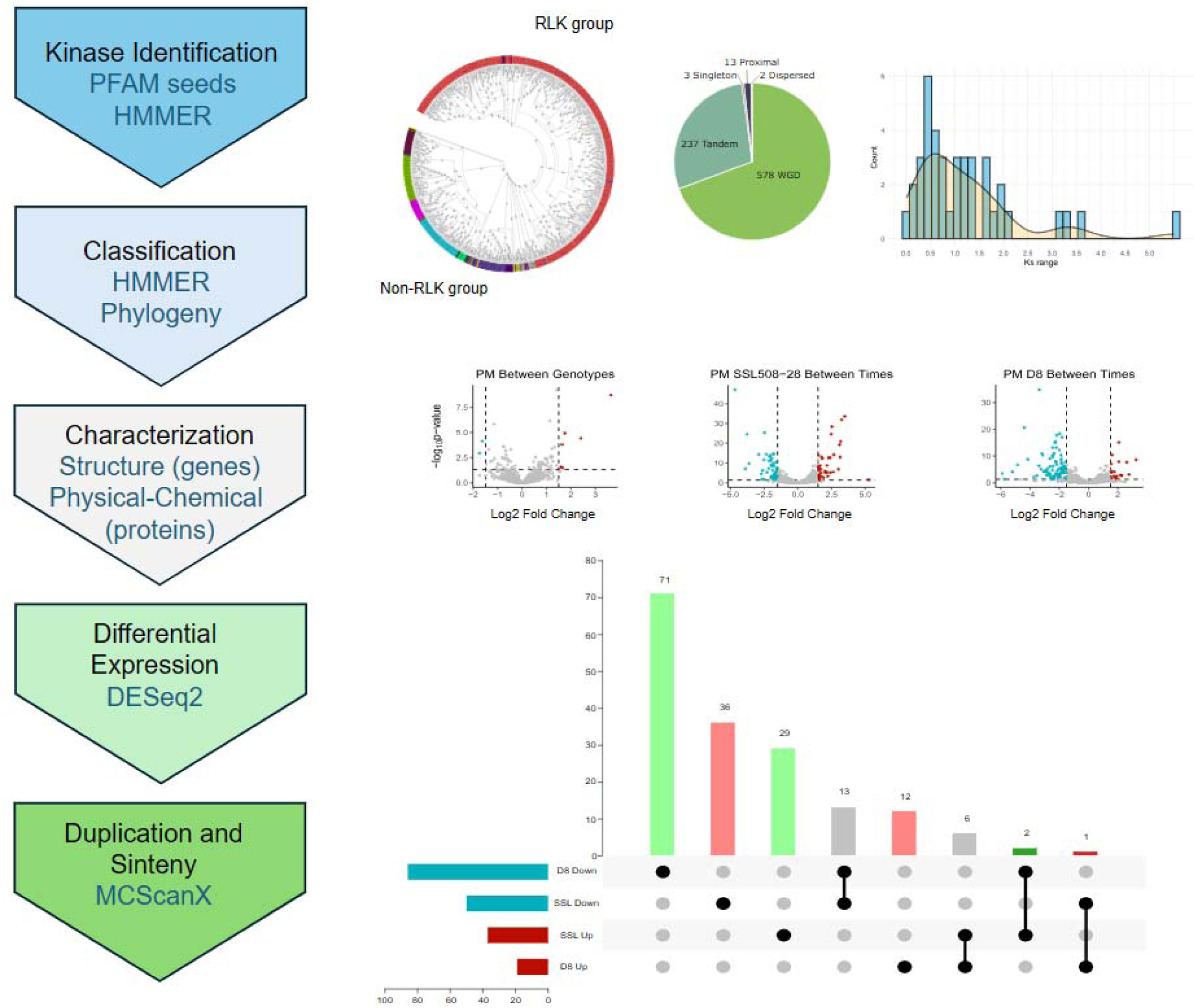

## 1. Introduction

Cucumber (*Cucumis sativus* L.) is a species that belongs to the *Cucurbitaceae* family, with significant importance for several industries worldwide.^1^ This crop is prominent in large and small-scale agricultural systems, contributing substantially to the economy. The demand for cucumbers, whether in their fresh or processed form, boosts agricultural production and strengthens the development of international trade alliances, ensuring year-round availability. Cucumbers are essential to the food industry due to their nutritional potential and medicinal attributes, mainly their high potassium content, which contributes to the relief of blood pressure.^1,2^ Given its applicability, cucumber cultivation has emerged as an economically viable alternative for several market sectors, including the production of plant-derived pharmaceuticals.^3^ Furthermore, within the beauty and skincare industry, cucumbers are recognized for their moisturizing, refreshing, and calming properties.^4^

Despite its agronomic relevance, the susceptibility of cucumbers to diseases and pests results in a reduction in foliage, consequently leading to a decrease in productivity.^5^ Powdery mildew (PM), caused by fungal pathogens such as *Podosphaera xanthii* (Castagne) U. Braun & Shishkoff (syn. *Sphaerotheca fuliginea* (Schlechtend.:Fr.) Polacci) and *Erysiphe cichoracearum* DC. (syn. *Golovinomyces cichoracearum*),^6,7^ represents a significant threat. According to Chen et al.,^8^ the main families of genes that confer resistance to PM are MLO (Mildew Locus O),^9^ PMR (Powdery Mildew Resistance),^10^ and TCTP (Translationally Controlled Tumor Protein).^11^ The cucumber genome encodes 14 MLO genes, with three of them grouped in Clade V (CsMLO05, CsMLO13, and CsMLO14),^12^ which contains the known PM susceptibility genes to PMs in other dicots. One of these genes, CsaMLO1, co-localizes with a previously identified QTL for PM resistance and its expression was upregulated after inoculation with *P. xanthii*.^13^ Additionally, homologs of the PMR4 and PMR5 susceptibility genes have also been found in the cucumber genome.^14^ Progress in PKs research links the LRR-RLK (Leucine-Rich Repeat Receptor-Like Kinase) as R genes for PM,^8^ along with the CRK (Cysteine-Rich Receptor-Like Kinase) family identified in key effective QTL for resistance to PM.^15^

It should be noted that many advances were achieved after the publication of the cucumber genome,^16^ including the possibility of studying gene superfamilies, such as protein kinases.^17^ Although a comprehensive characterization of the cucumber kinome remains limited, specific PK families have received attention in genomic analyses, contributing significantly to our understanding of their functions and potential involvement in various biological processes and stress responses. It is worth noting that several genomic analyses have already been conducted focusing on specific PK families within this species. To illustrate, the Casein Kinase (CK) family, which includes proteins responsible for phosphorylating casein,^18^ plays roles related to the circadian cycle,^19^ plant growth, and the regulation of light-modulated gene expression.^20^ On the other hand, the Mitogen-Activated Protein Kinase (MAPK) family constitutes a group of serine/threonine kinases that participate in diverse signal transduction pathways associated with hormonal responses and substantial developmental changes in organisms.^21^ Conversely, the Lectin Receptor-Like Kinase (LecRLK) family emerges as notably crucial in the innate immunity of plants, especially in defense against biotrophic pathogen, such as *P. xanthii*, the causal agent of PM.^22^ Furthermore, the Calcium-Dependent Protein Kinase (CDPK) and CDPK-Related Protein Kinase (CRK) families play roles in calcium-dependent signaling pathways in response to environmental stresses.^23,24^ PKs play a key role in modulating cellular responses to external stimuli,^25^ highlighting their importance in enhancing cucumber genetic improvement. Primarily, PKs’ main function is cellular signaling and regulation by phosphorylating PKs’ targets, a critical biological process that enables signaling within gene expression networks in various cellular processes.^26^ Thus, the comprehensive characterization of a kinome and its analysis holds substantial importance, particularly in the context of plant immunity.

Given the complexity and the need for a deeper understanding of the cucumber kinome, this study adopts an exploratory research approach. The results of this work enabled us to comprehensively evaluate the structural and evolutionary genomic attributes of PK genes and their expression in the context of PM. By focusing on a broad exploration of gene expression patterns under PM stress, we aim to generate fundamental knowledge that can pave the way for further hypothesis-driven research. This work addresses current gaps in the literature and provides a rich dataset for subsequent targeted studies that aim to increase disease resistance in cucumber and related species within the Cucurbitaceae family.

## 2. Material and Methods

### 2.1. Genome-wide identification and classification of cucumber PKs

The identification of *Cucumis sativus* PKs was based on the alignment of the Pkinase (PF00069) and Pkinase_Tyr (PF07714) subfamilies against the cucumber Gy14 annotated proteins, available in the Cucurbit Genomics Database v2 (CuGenDbv2).^27^ Such an alignment was performed with hidden Markov models (HMMs) obtained from the Pfam database (http://pfam.xfam.org/)^28^ together with the HMMER tool^29^ considering an E-value cutoff of 0.01.

After identifying the cucumber PKs, we classified them into subfamilies using the HMMER tool together with the family HMMs estimated from PKs of 25 plant species, as described and available as supplementary files by Lehti-Shiu et al.^17^: *Aquilegia coerulea* E. James*, Arabidopsis lyrate* (L.) O’Kane & Al-Shehbaz*, Arabidopsis thaliana* (L.) Heynh.*, Brachypodium distachyon* (L.) P.Beauv.*, Carica papaya* L.*, Citrus clementina* Hort.*, Citrus sinensis* L.*, Chlamydomonas reinhardtii* P.A. Dang.*, Cucumis sativus* L.*, Eucalyptus grandis* W.Hill*, Glycine max* (L.) Merr.*, Manihot esculenta* Crantz*, Medicago truncatula* Gaertn.*, Mimulus guttatus* (DC.) G.L.Nesom*, Oryza sativa* L.*, Populus trichocarpa* Torr. & A.Gray ex Hook.*, Prunus persica* (L.) Stokes*, Physcomitrella patens* (Hedw.) Mitt.*, Ricinus communis* L.*, Selaginella moellendorffii* Hieron.*, Setaria italica* (L.) P.Beauv.*, Sorghum bicolor* (L.) Moench.*, Vitis vinifera* L.*, Volvox carteri* F. Stein, and *Zea mays* L.

To confirm the obtained family classification, we estimated a phylogenetic tree of amino acids of PK sequences, which were aligned using the MAFFT v7.453 program.^30^ The phylogenetic tree was estimated with the FastTree 2.1.11 software,^31^ considering 1,000 bootstrap replicates and an approximately-maximum-likelihood approach together with the Whelan Goldman (WAG) model for amino acid evolution.^32^ As an outgroup, we selected a sequence from the *Chlamydomonas reinhardtii* v5.6 proteome (Cre07.g349540.t1.1) retrieved from Phytozome (https://phytozome-next.jgi.doe.gov).^33^ We selected this species because it does not have a multicellular ancestor,^34^ which indicates that it must have evolved before cucumber proteins.

### 2.2. Characterization of the cucumber PK sequences

The evaluation of gene structure of cucumber PKs was performed considering the number of exons and introns retrieved from the Gy14 GFF file, obtained from the CuGenDBv2.^27^ The chromosomal positions of PK genes were also obtained from the Gy14 GFF file, and a physical map of the chromosomes was constructed using the TBtools software.^35^ The analysis of the composition of the PK domains was performed using the Pfam database and the HMMER tool.^29^ For the physical and chemical characterization of PKs, we estimated using the ProtParam module from the SeqUtils subpackage:^36^ (i) the isoelectric point; (ii) the molecular weight; and (iii) the general average hydropathy (GRAVY). For the number of amino acids (aa), predictions of transmembrane helices, and signaling peptides, the TOPCONS server (https://topcons.cbr.su.se/pred/) was employed.^37^ We estimated the potential subcellular locations of PKs using the BUSCA web server (http://busca.biocomp.unibo.it/).^38^ Finally, for functional annotation, we used the Blast2GO^39^ tool and retrieved the associated Gene Ontology (GO) terms to create a treemap using the REViGO tool (http://revigo.irb.hr/)^40^ along with the treemap R package.^41^

### 2.3. Kinase gene expression patterns associated with Powdery Mildew

Initially, we assessed the expression of PK genes using raw RNA sequencing (RNA-Seq) data derived from an independent experiment retrieved from the CuGenDBv2 database (Table 1). This dataset was chosen based on its direct association with the response of cucumber to biotic stress caused by *P. xanthii*.^24^ After fitting the model with the full set of genes associated with Powdery Mildew, the subset differentially expressed PKs was enriched for further analysis.

**Table 1.** Description of the data considered for the cucumber kinome construction.

| Condition | Pathogen lifestyle | Resistant | Susceptible | RNA<br>extraction | Data origin |
| --- | --- | --- | --- | --- | --- |
| PM | Biotrophic | SSL508-28 | D8 | 0 × 48 h | Xu et al. <sup>24</sup> |

PM: powdery mildew (*Podosphaera xanthii*), h: hours post-inoculation.

To evaluate resistance to PM, experimental trials were carried out during three consecutive years (spring 2013, spring 2014 and autumn 2015) on the SSL508-28 (resistant) and D8 (susceptible) genotypes, under greenhouse conditions (Yangzhou, China). Inoculations were performed by applying conidia collected from naturally infected D8 plants, as described by Xu et al.^24^ A total of 12 standard Illumina paired cDNA libraries were prepared, including three biological replicates for each genotype evaluated at 0-and 48-hours post-inoculation. These libraries were sequenced using the Illumina HiSeq 2500 platform, producing 125 base pair paired-end reads. For mapping the reads to the cucumber reference genome, high-quality nucleotides were filtered according to their quality score (Q > 20) and aligned to the 9930 (Chinese Long) v2 reference genome^42^ using TopHat v2.0.9.^43^ To count the number of reads mapped to reference genes, Cufflinks v2.1.1^44^ was employed. The raw RNA-seq data was generated and available by Xu et al.^24^

To identify the whole set of DEGs, we employed the raw count data as input for DESeq2 package.^45^ We considered the following contrasts: PM between genotypes (time as covariate); D8 between time points (0h vs 48h); and, SSL508-28 between time points (0h vs 48h). For each comparison, we conducted negative binomial tests to identify DEGs, with a significance threshold set at a maximum adjusted p-value (false discovery rate, FDR) of 0.05 and a minimum absolute log2 fold change (log2fc) of 1.5. After filtering the PK-related DEGs, they were visualized using Volcano Plots, generated using the tidyverse,^46^ RColorBrewer,^47^ and ggrepel^48^ packages in R (https://www.r-project.org/).^49^

For a better comprehension of the families that respond similarly to each disease, we employed Venn diagrams, using TBtools^35^ to illustrate the overlap between the total PK DEGs and those that were either up-regulated and down-regulated in the resistant and susceptible genotypes. The integration of this information allowed us to indicate genes potentially related to resistance or susceptibility.

After determining the DEGs, we began an approach to identify their influence on the plant in response to disease through a comparative analysis to identify the profile of R-related (resistant) and S-related (susceptible) genes. Putative R-genes were thus assigned if they were upregulated in the resistant genotype or downregulated in the susceptible genotype. In the same way, the putative S-genes are either upregulated in the susceptible genotype or downregulated in the resistant genotype. The other genes that did not fit this profile were disregarded from the analysis.

### 2.4. Kinase duplication and synteny analyses

To conduct duplication analysis and identify categories of duplication events, we employed the Multiple Collinearity Scan toolkit (MCScanX).^50^ Calculations of synonymous (Ks) and non-synonymous (Ka) substitution rates were performed using the MCScanX module hosted in the TBtools software.^35^ The time elapsed for each duplication event was determined using the formula T = Ks/2λ, where λ represents the mean synonymous substitution rate (6.5 x 10^-9^).^51^

A comprehensive synteny analysis was performed to explore the relationships among kinase genes in the Gy14 cucumber and DHL92 melon^52^ genomes, both sourced from the CuGenDBv2 database. The identification of syntenic blocks was accomplished through the application of the MCScanX toolkit, while subsequent visualization of the data was achieved using the Dual Synteny Plot package, seamlessly integrated into the TBtools software. This methodological approach allowed a detailed examination of conserved genomic regions and facilitated a visually informative representation of the syntenic relationships between kinase genes in the cucumber and melon genomes.

#### 2.4.1. Safety Information

This work was conducted entirely through in silico methods using publicly available datasets. No experimental work involving hazardous chemicals, materials, or biological agents was performed during the course of this research.

## 3. Results

### 3.1. Genome-wide identification and Classification of cucumber PKs

Alignment of the annotated set of cucumber Gy14 proteins against two HMM profiles, Pkinase (PF00069) and Pkinase_Tyr (PF07714) identified 837 putative PKs, and after a domain check two atypical PKs were removed, remaining 835 true PKs. According to the best hit, 517 proteins presented the Pkinase domain (PF00069), while 318 presented Pkinase_Tyr (PF07714) (Supplementary Tables S1). Analysis of the domain composition of the 835 PKs in our study showed that there are 5,930 domains, of 558 different types (Supplementary S2). The average number of domains per protein was seven, with a wide variation, from 1 to 22 (Supplementary Table S3). A total of 833 PKs were predicted to have the PKinase domain (PF00069), while 824 were predicted to have the Pkinase_Tyr domain (PF07714). Interestingly, most PKs (822) contained both domains, with only 13 PKs having one or the other. Besides the PKinase and Pkinase_Tyr domains, the most frequent additional domains included APH (348 PKs), Kinase-like (297), Pkinase_fungal (243), LRR_1 (192), LRR_4 (192), LRR_8 (192), Haspin_kinase (188), Kdo (179), LRRNT_2 (142), ABC1 (134), and LRR_6 (131). Giving special emphasis to the different types of LRR domains, we identified LRR_1 in 192 proteins, as well as, LRR_4 (192), LRR_8 (192), LRRNT_2 (142), LRR_6 (131), LRR_9 (17), LRR_2 (7), LRR_5 (7), LRR_12 (1) and LRRCT (1). From an evolutionary perspective, it is important to highlight that numerous combinations have been detected between all these domains (Supplementary Table S3).

Domain composition analysis using the Pfam database and the HMMER tool identified two sequences as atypical PKs due to the absence of canonical kinase domains: (i) CsGy6G033750.2, which belongs to the ABC1 family (PF03109.16); and (ii) CsGy1G024940.2, associated with the Haspin_kinase family (PF12330.8). It is worth noting that atypical kinases, despite lacking canonical kinase domains, may still retain kinase functions, as reported in prior studies.^53,54^ However, as our study primarily focuses on typical kinases, we have removed these atypical kinases from further investigations.

Based on the HMM approach proposed by Lehti-Shiu et al.,^17^ we were able to classify the 835 genes encoding cucumber kinases into 20 distinct groups and 123 families (Figure 1 and Supplementary Figure S1, Tables S4 and S5), namely: AGC (cAMP-dependent protein kinases, cGMP-dependent protein kinases, various types of protein kinase C, protein kinase B, 3-phosphoinositide-dependent protein kinase-1, and the ribosomal protein S6 kinases), Aur (aurora kinase), BUB (budding uninhibited by benzimidazoles), CAMK (calcium/calmodulin-dependent protein kinase), CK1 (casein kinase 1), CMGC (cyclin-dependent kinase, mitogen-activated protein kinase, glycogen synthase kinase, and cyclin-dependent-like kinase families), Plant specific (Group-Pl), IRE1 (inositol-requiring enzyme 1), NAK (NF-κB-activating kinase), NEK (never in mitosis gene-A), PEK (pancreatic eukaryotic initiation factor 2 α-subunit kinase), RLK (receptor-like kinase), SCY (*Saccharomyces cerevisiae* [yeast] kinase), STE (serine/threonine kinase), TKL (tyrosine kinase-like kinase); TLK (tousled-like kinase), TTK (threonine/tyrosine kinase), ULK (unc-51-like kinase), WEE (wee1, wee2, and myt1 kinases), and WNK (with no lysine-K).

**Figure 1.**
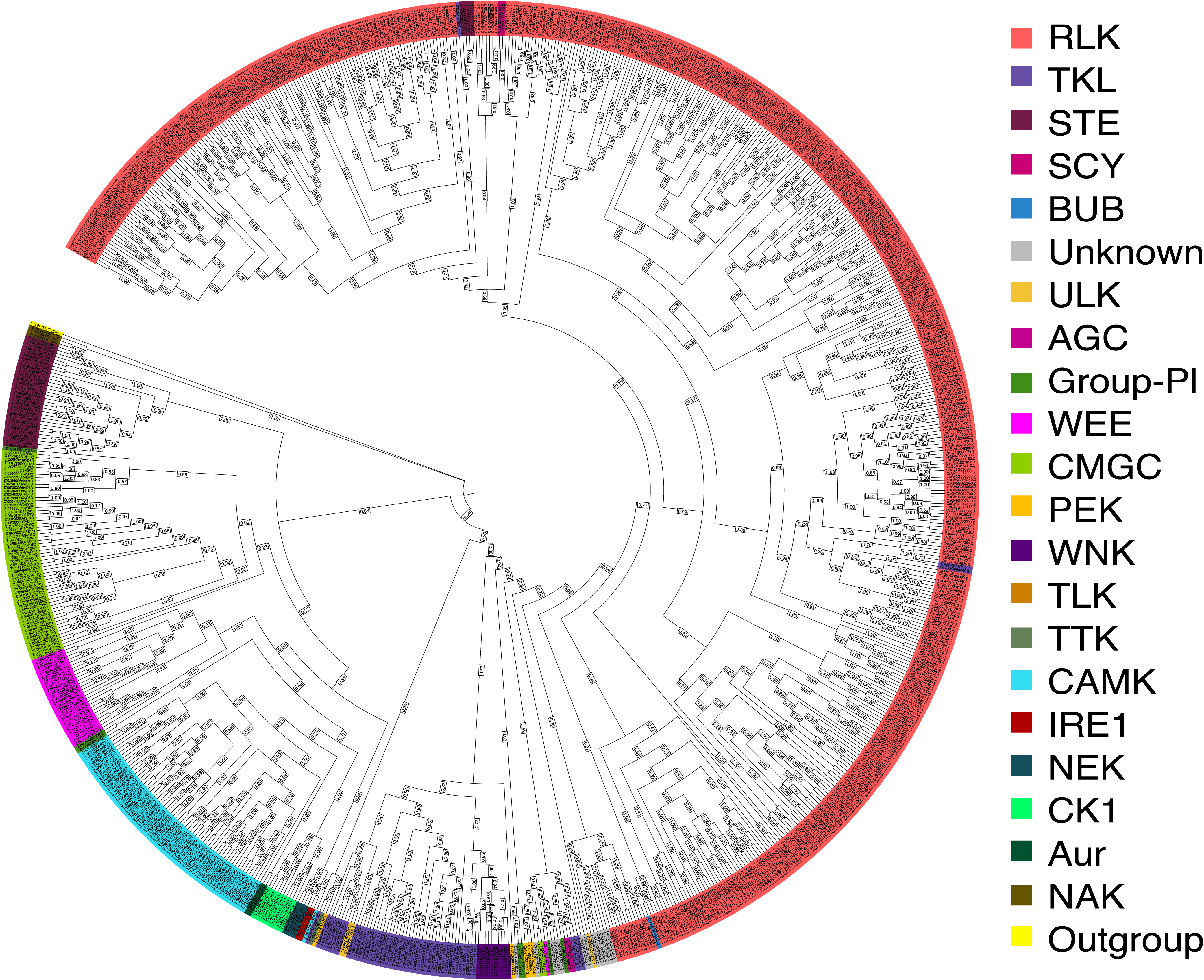
Phylogenetic tree of *Cucumis sativus* PKs with the approximately-maximum-likelihood Likelihood approach with bootstrap values (> 70%), using the Whelan Goldman (WAG) model with gamma parameter. The clades highlighted are referred to the groups RLK, SCY, TKL, STE, CMGC, AGC1, CK1, and CAMK (from up to down). The members of each family are collapsed to a better visualization. The gene names displayed with square brackets represent branches with single genes. The tree is rooted with a kinase (Cre07.g349540.t1.1) from the *Chlamydomonas reinhardtii* v5.6 proteome. Some genes from expanding groups can be found within other consolidated groups.

To confirm the family classification, we used a phylogenetic approach that effectively grouped the proteins into eight distinct clusters with strong support, as indicated by the 70% bootstrap threshold. These clusters were related to the AGC, CAMK, CK1, CMGC, RLK, SCY, STE, and TKL groups (Figure 1, and Supplementary Figure 1). The remaining proteins exhibited substantial functional diversity and expansion, forming different groupings of families within the same group. Furthermore, 13 genes could not be classified within their respective group (in instances where there was only one gene per family) or within their respective families (in cases where a family possessed multiple genes). These unclassified genes were considered as belonging to an "unknown" category in the cladogram.

Among the groups defined for the cucumber kinome, the RLK group was the most numerous, with 530 genes (63.5% of the total number of PKs) according to HMMER and 527 confirmed by phylogeny analysis, with exceptions of CsGy1G009250.2, CsGy1G031510.2, and CsGy6G033380.1. The genes of this group were subdivided into 57 families according to HMMER and into 56 families based on phylogenetic analysis. Notably, 35 of these families contained only one gene, while the RLK-Pelle_DLSV family was the most abundant with 68 genes, followed by the RLK-Pelle_LRR-XI-1 (40 genes), RLK-Pelle_LRR-III (38 genes), RLK-Pelle_RLCK-VIIa-2 (32 genes), and RLK-Pelle_L-LEC (28 genes) (Supplementary Tables S4 and S5).

### 3.2. Characterization of the cucumber PK sequences

The distribution of the 835 PK genes across chromosomes reveal a specific pattern of gene localization in this superfamily (Figure 2), for the accumulation of PKs in the telomeric regions, aligning with a tendency for resistance genes to be in the distal regions of telomeres.^55^ Additionally, genes from the same family tended to cluster together, suggesting potential tandem duplications. Chromosome 3 had the highest number of PK genes (170) and chromosome 7 had 82. It is noteworthy that two genes were associated with cucumber scaffolds (chromosome 0) and were not allocated to any chromosome in the Gy14 genome (Figure 3A).

**Figure 2.**
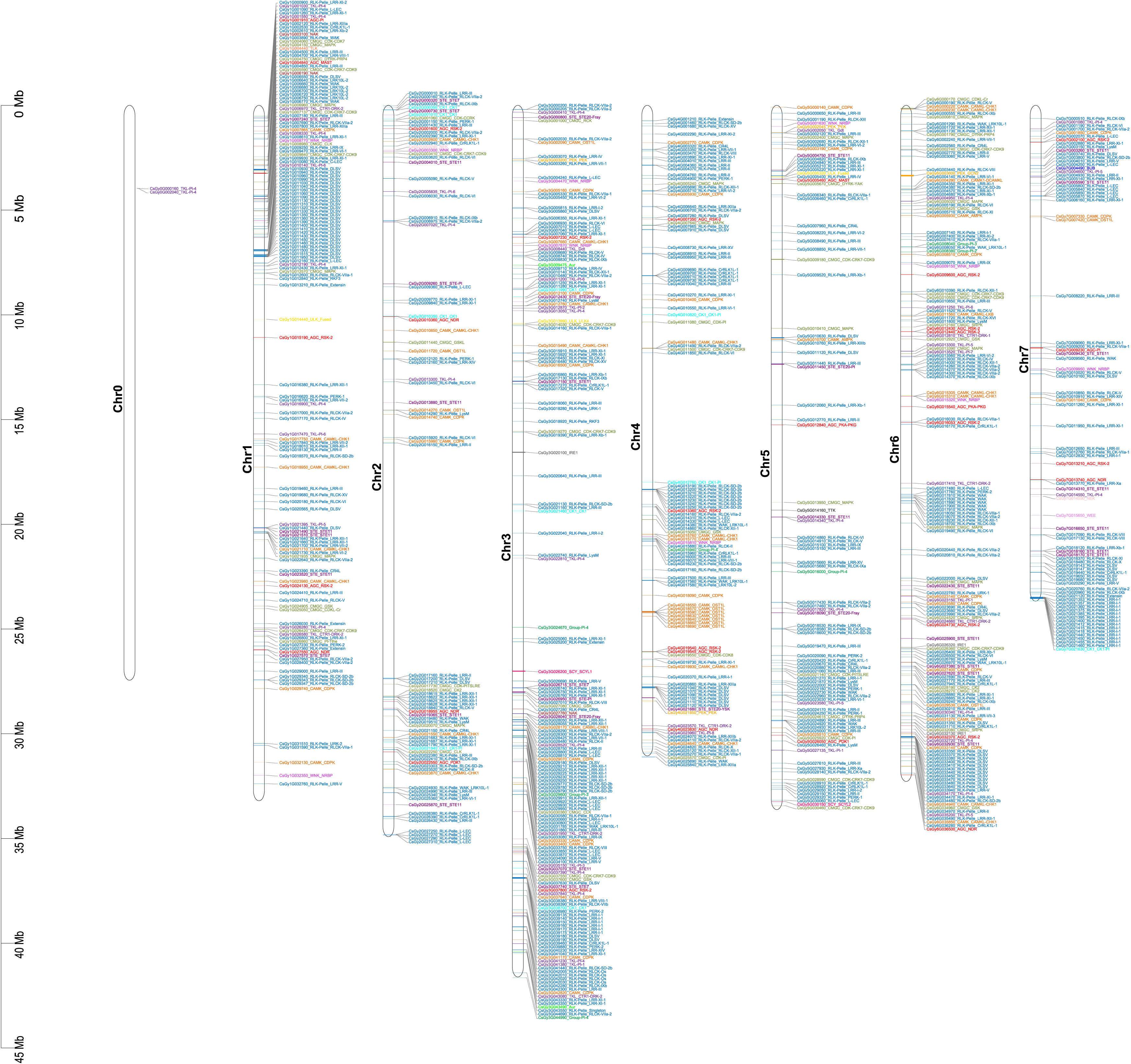
Chromosome physical localization of kinase genes from *Cucumis sativus*. The kinase genes are presented by their names followed by the family name, and the colors represent different groups. Chromosome zero (0) contains genes that have not been assigned to specific chromosomes.

**Figure 3.**
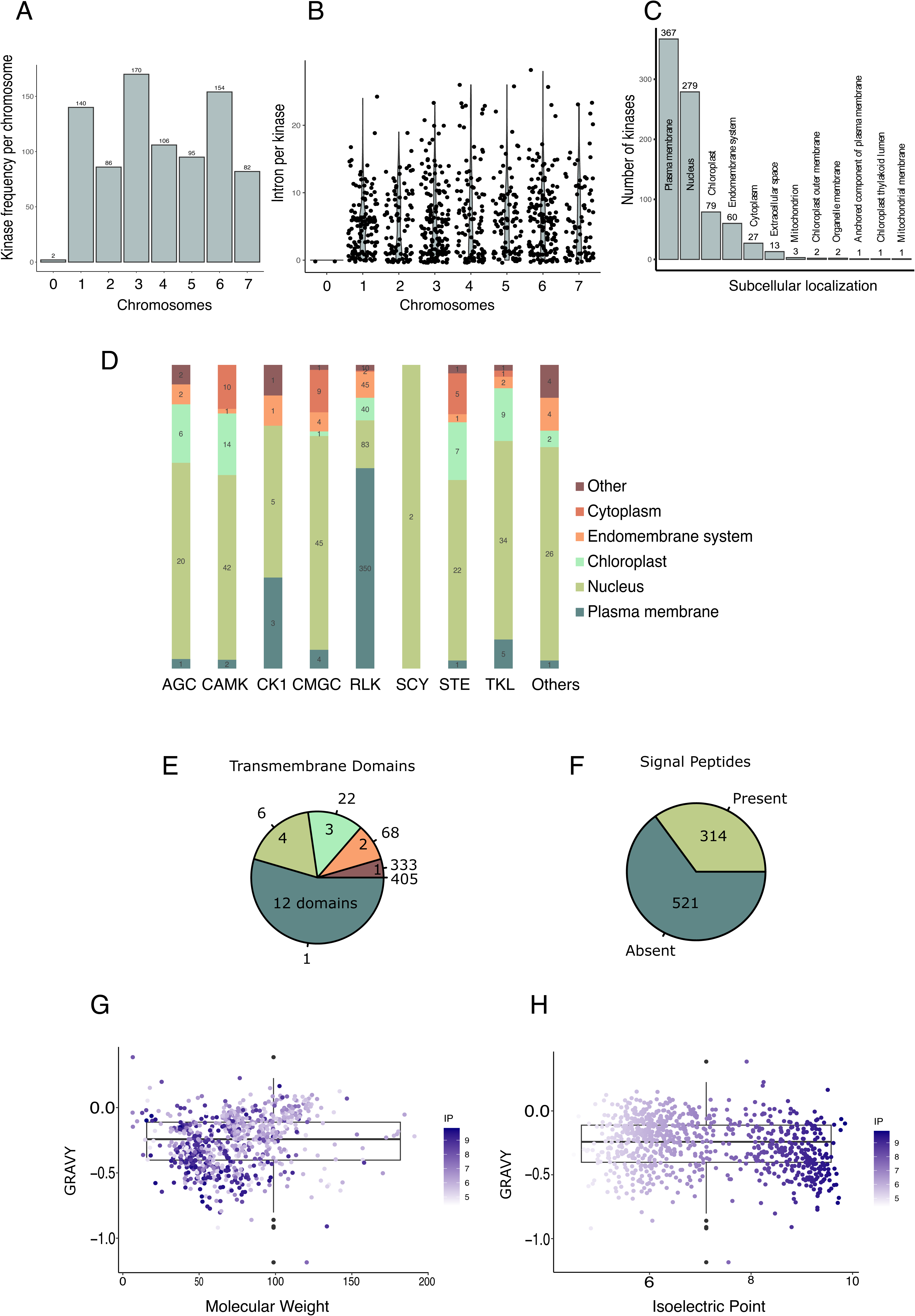
**(A)** Number of cucumber kinases per chromosome. **(B)** Number of introns per kinase per chromosome. **(C)** Number of kinases per subcellular location. **(D)** Distribution of kinase per subcellular location per kinase group. **(E)** Number of transmembrane domains and signal peptides. **(F)** molecular weight (MW), and isoelectric point (IP) in function of grand average of hydropathy (GRAVY).

Analyzing the structure of the 835 genes encoding cucumber PKs, we observed a variable number of introns, from 0 to 28, with a median of 5 introns per gene (standard deviation of 5.46). The highest number of introns was observed in the CsGy6G003600 gene, a member of the PEK_GCN2 family, which presented 28 introns. Among the total of 835 PK genes, 123 (14.7%) presented no introns, while 554 (66.3%) contained between 1 and 10 introns; 139 (16.7%) had between 11 and 20 introns, and 19 (2.3%) exhibited a substantial range of 21 to 28 introns (Figure 3B, Supplementary Table S6).

The subcellular location prediction results revealed that approximately 44% of PKs are potentially located in the plasma membrane, 33% in the nucleus, 9% in the chloroplast, 7% in the endomembrane system, 3% in the cytoplasm, and 2% in the extracellular space (Figure 3C). Intriguingly, the different groups of protein kinases present a particular subcellular localization profile, with important similarities and highlights, such as the fact that a significant portion of RLKs are positioned in the plasma membrane. The prediction of a significant number of PKs for the nucleus is also worth highlighting (Figure 3D; Supplementary Table S7).

The number of transmembrane (TM) domains per kinase exhibited significant variation. Among the highlights, we found that they are not present in 405 PKs; 333 PKs present one, while there is one PK (CsGy6G023990.2) with 12 TM domains (Figure 3E; Supplementary Table S7). Among the 835 PKs, 314 presented a signal peptide, which provides more information about the initial fate of these proteins. (Figure 3F; Supplementary Table S7). It is important to reflect on the presence of transmembrane domains and signal peptides in proteins, since this analysis does not only include proteins located in the plasma membrane, but also those acting in other internal membranes. The presence of a signal peptide, in turn, suggests action dependent on the secretory pathway, whether in the transmembrane or extracellular domain.

The physicochemical characteristics of the PKs were highly variable (Figure 3G-H and Supplementary Table S7). The number of amino acids ranged from 57 to 1694 (median: 645); molecular weight spanned from 6.42 to 191.01 kDa (median: 71.27 kDa); the isoelectric point ranged from 4.38 to 9.85 (median: 6.57); and the hydropathy (GRAVY) varied from -1.19 to 0.39 (median: -0.24). This diversity in physical and chemical properties highlights the wide range of sizes, charges, and hydrophobicity characteristics of the PKs, providing insights into the intricate mechanisms by which proteins harness a highly complex enzymatic system to modulate responses to environmental stress.

To corroborate the functional diversity of this superfamily, the functional annotation revealed a total of 386 different GO terms occurring 4,273 times. These GO categories encompass molecular function (F) (95 terms), biological processes (P) (227 terms), and cellular components (C) (64 terms). These annotations provide valuable insights into the functions and locations of PKs, highlighting their involvement in a wide range of metabolic processes and cellular components. The Gene Ontology (GO) classification of cucumber PKs also provides insightful information about the essential roles they play. Within the scope of Metabolic Processes (P), protein phosphorylation (667 terms), peptidyl-tyrosine phosphorylation (63), and signal transduction (62) presented the highest numbers of terms (Figure 4A-B). Coherently, among the Molecular Functions (F), ATP binding (624) and kinase activities such as protein serine/threonine kinase activity (409 terms) are the most numerous. Regarding cellular components (C), the action of PKs in the context of the plasma membrane is endorsed by the largest number of terms (272 and 174) (Supplementary Table S8).

**Figure 4.**
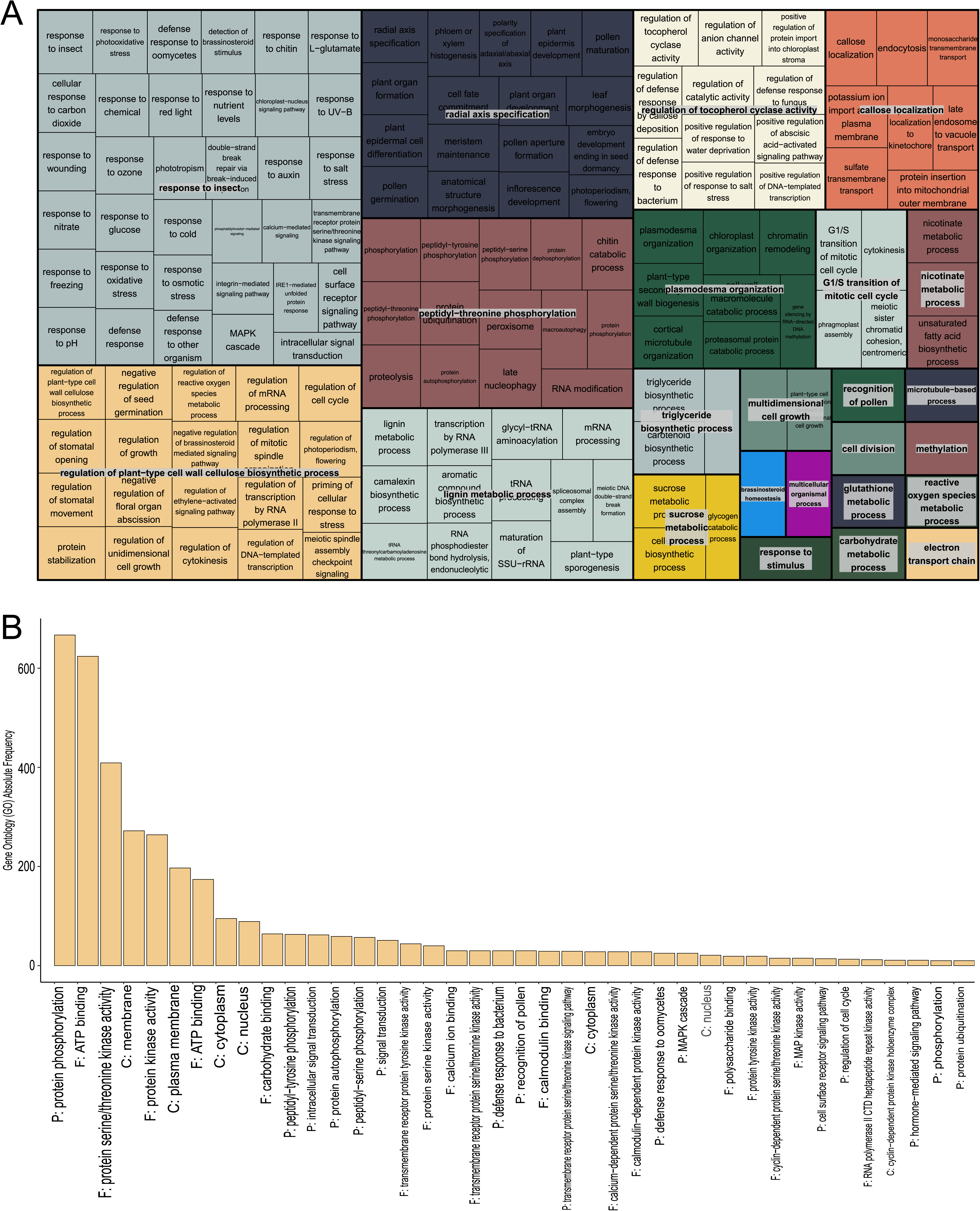
**(A)** Biological processes (227) predicted in the Gene Ontology analysis. **(B)** P: biological process, F: molecular function, C: cellular component. Figure shows the 40 top GO terms; a full list is available in Supplementary Table S7.

### 3.3. Kinase gene expression patterns associated with Powdery Mildew

We evaluated the involvement of cucumber PKs in response to the powdery mildew pathogen in a transcriptome of this pathosystem obtained by Xu et al.^24^ To identify DEGs among the 22,626 genes, with enrichment for the 835 protein kinases in our study, we analyzed three contrasts, as detailed in table 2: (i) a comparison between resistant and susceptible genotypes while incorporating time as a covariate; (ii) evaluation of changes in gene expression in the resistant genotype after 48 hpi relative to time 0 hpi; and, (iii) an evaluation of gene expression alterations in susceptible genotypes after 48 hpi relative to time 0 hpi.

**Table 2.** DEG analysis of the compatible and incompatible interactions of cucumber genotypes against the pathogenic fungus *P. xanthii* (PM).

| Condition | Contrast | Contrast<br>time | Genes for<br>modeling | DEGs | PK DEGs | Down | Up |
| --- | --- | --- | --- | --- | --- | --- | --- |
| PM | SSL508-28<sup>R</sup><br>× D8<sup>S</sup> | covariate | 17,839 | 319 | 8 | 2 | 6 |
|  | SSL508-28<br>× SSL508-28 | 0h × 48h | 17,522 | 1,428 | 87 | 50 | 37 |
|  | D8 × D8 | 0h × 48h | 18,314 | 1,690 | 105 | 86 | 19 |

DEG: differentially expressed genes, PK: protein kinase, PM: powdery mildew (*P. xanthii*), (R): resistant, (S): susceptible, dpi: days post-inoculation. Down: down-regulated, Up: up-regulated.

For each comparison, we estimated a distinct model using the DESeq2 package and retained different numbers of genes for differential expression evaluation, ranging from 17,839 to 18,314 (Table 2). Following this, we identified DEGs, considering the whole number of genes in the species and subsequently narrowed our focus to the PK DEGs for further in-depth analyses. The number of DEGs varied from 319 to 1,690, while PK DEGs numbered from 8 to 105. In general, we observed a higher number of DEGs in the susceptible genotype. When considering time as a covariate, i.e., contrasting the gene expression between genotypes, we identified 319 DEGs (including 8 PKs) for the PM pathosystem. These DEG numbers were lower in comparison to the other contrasts performed.

Among the genotypes, only 8 significant PK differentially expressed genes were identified, with 2 being down-regulated and 6 up-regulated in the resistant genotype (Table 2; Figure 5; Supplementary Table S9). Considering the time factor for each cultivar, a total of 87 significant PK DEGs were observed in the resistant genotype, being 50 down-regulated and 37 up-regulated. In the susceptible genotype, 105 significant PK DEGs were identified, being 86 down-regulated and 19 up-regulated. When using the response time as a covariate, only 8 PK genes differentially expressed were observed in the PM pathosystem, indicating that these genes are consistently activated or silenced along with the time (0 or 48 hours), with the level of expression increasing or decreasing over time. Among these DEGs, six were activated in the resistant genotype, suggesting their significant roles in defense against PM: CsGy1G023520 (STE_STE11/chloroplast), CsGy2G025870 (STE_STE11/nucleus), CsGy7G003260 (STE_STE11/chloroplast), CsGy6G022000 (RLK-Pelle_DLSV/plasma membrane), CsGy6G034480 (RLK-Pelle_RLCK-SD-2b/endomembrane system), and CsGy7G002100 (AGC_RSK-2/extracellular space).

**Figure 5.**
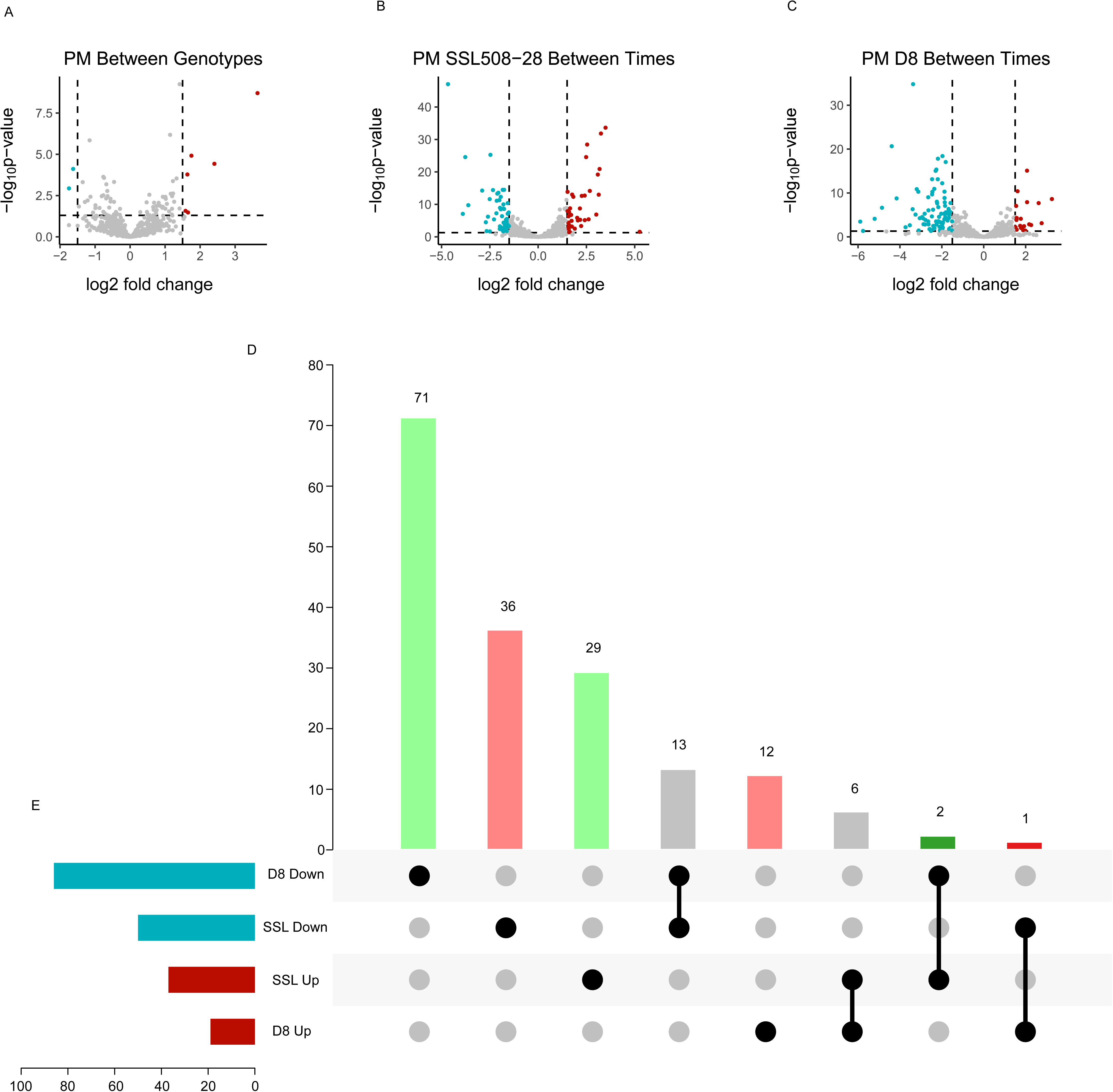
Differential expression of kinase genes (kinase DEGs): **(A)** Between genotypes (time as a covariate), **(B)** resistant (SSL508-28) and **(C)** susceptible (D8) genotypes to powdery mildew (PM) (*Podosphaera xanthii*). **(D)** Putative R-genes (light green), Putative S-genes (light red), Like-R genes (dark green), Like-S genes (dark red), Unrelated genes (gray). **(E)** Down-regulated genes in resistant and susceptible genotypes (blue), and up-regulated genes in resistant and susceptible genotypes (red). SSL: SSl508-28.

It was observed that the number of up-regulated genes was lower than that of down-regulated genes in both genotypes due the time. Among the observations, we highlight six genes of particular importance due to their high levels of expression (Log2FC ranging from 3.09 to 5.25): CsGy6G005190 (RLK-Pelle_RLCK-VI/nucleus), CsGy1G021935 (CMGC_MAPK/cytoplasm), CsGy7G005800 (RLK-Pelle_L-LEC/nucleus), CsGy6G002560 (RLK-Pelle_CR4L/plasma membrane/also activated in the susceptible -D8), CsGy3G012100 (CAMK_CDPK/nucleus/also activated in the susceptible - D8), CsGy1G027950 (RLK-Pelle_RLCK-VIIa-2/nucleus/also activated in the susceptible - D8). The first three PK genes listed had their activity silenced in the susceptible genotype (D8), suggesting that their activity may be directly involved in defense against PM. In the susceptible genotype (D8), it is noteworthy that the genes with the lowest expression levels were all down-regulated. This observation suggests that gene silencing through various mechanisms can cause susceptibility, but this hypothesis needs further investigation to identify the mechanisms responsible for gene silencing.

According to previously established criteria to consider genes related to resistance and susceptibility (putative R-genes and S-genes), it was possible to perceive a spectrum of possibilities. There are genes that are upregulated only by the resistant genotype SSL508-28 (putative R-genes), as well as by the susceptible genotype D8 (putative S-genes). This same scenario was verified considering the downregulated genes. Genes upregulated by resistant genotypes and negatively regulated by susceptible genotypes were considered as candidate genes for resistance functions (like R-genes), since they may have made a positive difference in preventing the emergence of diseases in resistant genotypes or were missing for susceptible genotypes. This understanding is obtained when a resistance gene is transiently silenced.^56^ According to this view, genes upregulated in susceptible genotypes and negatively regulated in resistant genotypes can be considered candidates for susceptibility functions (like S-genes), since they may have contributed to the success of the pathogen or manifestation of symptoms by susceptible genotypes and ceased to perform these functions in the resistant genotype.^57^ From this set, genes up-regulated in both genotypes or down-regulated in both genotypes can be considered unrelated to the resistance or susceptibility phenotypes (Figure 5).

Among the putative R genes (upregulated in the resistant genotype or downregulated in the susceptible genotype), there was a predominance of RLK-Pelle receptor kinases, especially from the RLK-Pelle_DLSV and RLK-Pelle_LRR-XI-1 families. Genes associated with signaling cascades, such as members of the MAPK and CAMK families, were also identified as DEGs, suggesting an important role of signal transduction in the defense system. It is consistent to note that the analysis of subcellular localization showed that most of the candidate resistance genes are associated with the plasma membrane, as well as the nucleus, which signals the cellular perception of stress by the plant, followed by regulation of gene expression. Among the candidate susceptibility genes, this scenario is also notable.^58^ Some DEGs, classified as RLK-Pelle_LRR-I-1, were also associated with susceptibility, suggesting the possibility of divergent functions. In terms of subcellular localization, there is again a relative occurrence of plasma membrane and nuclear proteins, with some located in intracellular compartments such as the chloroplast and endomembrane system.

### 3.4. Kinase duplication and synteny analyses

Through an in-depth examination of duplication events, our study yielded estimations for 833 protein kinases (PKs), considering that two kinases are still discernible at the scaffold level in the genome. The predominant origin of these PKs was attributed to whole genome duplication (WGD) events, encompassing approximately 69.4% (578 PKs) (Figure 6A). Tandem duplications contributed to ∼28.5% (237 PKs), proximal duplications accounted for ∼1.6% (13 PKs), singleton events represented ∼0.36% (3 PKs), and dispersed duplications were observed at a minor frequency of 0.24% (2 PKs). For a subset of 40 genes, we calculated both the Ka and Ks substitution rates (Figure 6B). The Ka/Ks ratio, indicative of the interplay between purifying selection, neutral mutations, and advantageous mutations, ranged from 0.146 to 2.198, with a mean ratio of 0.146 (Supplementary Table S10). Ratios below 1 suggest selective pressure, values around 1 denote neutral effects, while ratios exceeding 1 signify advantageous mutations. Furthermore, the estimated ages of these duplication events were determined based on the Ks rate, spanning from ∼50.42 to ∼422.81 million years ago (MYA). This temporal insight provides a comprehensive perspective on the evolutionary dynamics of the PK landscape, shedding light on the selective forces and mutational processes that have shaped this vital component of the genome over extensive periods.

**Figure 6.**
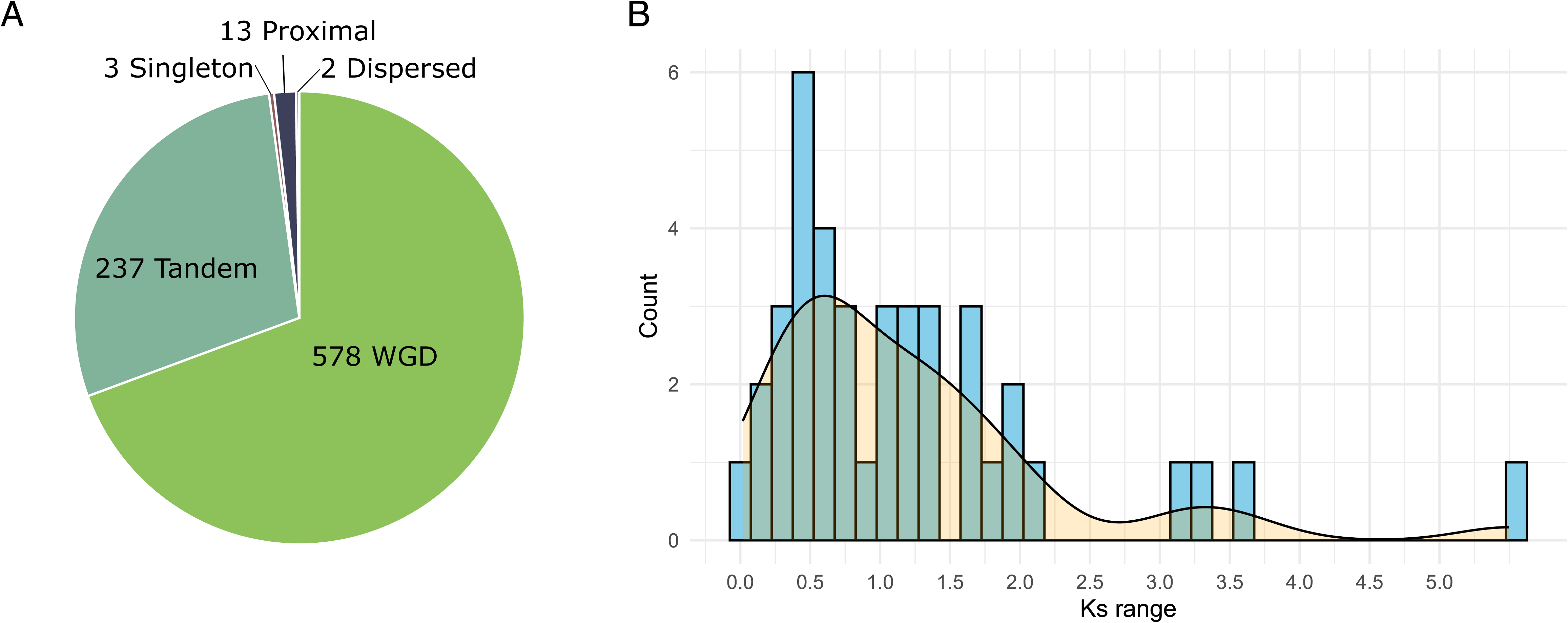
**(A)** Classification of duplication types of 833 Protein Kinases of *Cucumis sativus* (Gy14). **(B)** Range of Ks values for the Protein Kinases of *Cucumis sativus* (Gy14).

A total of 237 PKs exhibiting tandem duplications were identified across 54 families. Among these, one family belonged to the CAMK group, two to CMGC, 43 to RLK, 5 to TKL, and singular representatives from TLK, WEE, and WNK. Tandem duplications manifest when a chromosomal segment is replicated and integrated adjacent to the original segment, a result of the crossing-over process between chromatids.^59,60^ This mechanism plays a crucial role in amplifying the count of genes sharing identical or similar functions, contributing significantly to genomic diversity and functional specialization.

To investigate the evolutionary relationships between cucumber and melon species, we conducted a detailed synteny analysis, focused on cucumber kinase genes. Of the 835 kinase genes identified in cucumber, we identified remarkable syntenic relationships for 670 of these genes on all chromosomes of the two species (Figure 7, Supplementary Table S11). This finding encompasses 821 syntenic relationships with melon genes, of which 11 genes showed three relationships, 129 genes showed two relationships, while the remaining genes showed a single relationship. Analyzing the melon genome in relation to the cucumber PK array, we identified that 672 genes exhibited syntenic relationships, providing a comprehensive view of the genetic interconnections between these two species. These results provide insights into evolution and genetic relationships, highlighting complex patterns of conservation and divergence. The richness of these syntenic relationships reveals the intricate web of genetic connections between cucumber and melon, indicating not only shared ancestry but also specific duplication and rearrangement events throughout their evolutionary histories.

**Figure 7.**
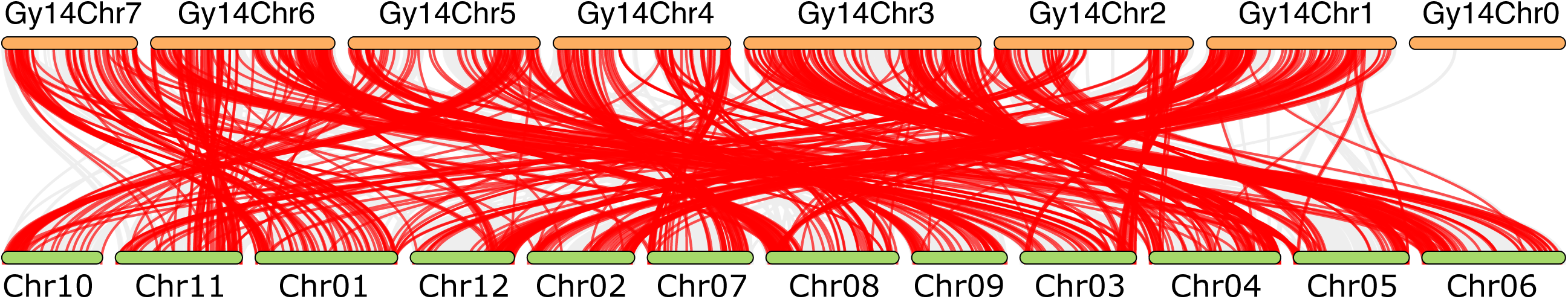
Synteny analysis of Protein Kinases between *Cucumis sativus* (Gy14) and *Cucumis melo* (DHL92). The chromosomes of *C. sativus* are represented in orange color and the prefix Gy14, and the chromosomes of *C. melo* represented in green and the prefix chr.

## 4. Discussion

### 4.1. Genome-wide identification and Classification of cucumber PKs

Cucumber was one of the first economically important crop to have its genome published. Over five years the genomes of the lineages 9930,^16,42^ PI 183967,^61–63^ Gy14 and B10 were made available,^27^ which provided a significant scientific advance in the field of cucumber genetic research. In addition, the utilization of recently developed high-throughput long-read sequencing technology has improved the quality of genome assembly for the Gy14 v2.1, 9930 v3.0, and B10 v3.0.^64^ These advances enabled us to perform a robust genome-wide analysis and obtain the first known cucumber kinome. Some projects have already been carried out considering some families of PKs in cucumber species,^17,65^ but a comprehensive superfamily analysis has been lacking. In this study, we performed bioinformatic analyses of the set of the cucumber annotated proteins to identify and characterize the PK superfamily.^66^ Using the profiles of Hidden Markov Models (HMM) PF00069 (Pkinase) and PF07714 (Pkinase_Tyr) to scan the cucumber annotated proteins, a total of 835 proteins containing canonical kinase domains were identified, which provided the opportunity for a holistic view of the biological processes played by PKs. Among these, 517 sequences were classified as Serine/Threonine kinase (Pkinase) and 318 sequences as Tyrosine kinase (Pkinase_Tyr). This number represents 3.69% of the total proteins of the cucumber species, slightly below the average (3.85%) of other species already studied, such as Arabidopsis,^67,68^ common bean,^69^ corn,^70^ cowpea,^71^ soybean^72^ and grapevine.^73^

For a better understanding of the evolutionary relationships between cucumber PKs, a phylogenetic analysis was obtained, which successfully grouped the members of eight well-established groups: AGC, CAMK, CK1, CMGC, RLK, SCY, STE and TKL.^17^ These groups consist of distinct families of PKs with known functions in several biological processes. However, there was significant functional diversity and expansion within these groups, as evidenced by the presence of different families within the groups. The expansion of these families may be related to the evolutionary process, since PKs are central regulators of stress in plants,^25^ acting like a CPU (Central Processor Unit), processing external signals and converting them into specific responses in the cell metabolism.^74^ As observed by Lehti-Shiu et al.^17^, some PKs of the remaining groups (AUR, BUB, PLANT-SPECIFIC, IRE1, NAK, NEK, PEK, TLK, TTK, ULK, UNKNOWN, WEE, and WNK) observed in our analysis are composed of PKs that probably arose after events that led to the divergence of plants, fungi and animals, with these families having few members, as they are absent in the algae *Chlamydomonas reinhardtii*, such as BUB, IRE1, PEK. This is an aspect that needs to be elucidated, regarding which families arose before and after the divergence of eukaryotes into fungi, plants and animals. Some specific families still have not their functions clarified in some plant species, such as the PKs of the Group-Pl^17^, NEK in both plants and animals.^75–77^

According to the rules proposed by Lehti-Shiu et al.^17^, the 835 PKs of the cucumber were classified into 20 groups. Out of these, the RLK group was the most numerous, comprising 527 genes, representing approximately 63.5% of the total kinases identified. This result is consistent with previous studies highlighting the abundance of RLKs in common bean, corn, Arabidopsis, vine, cowpea, and soybean plants, averaging 66.9%, where RLKs play crucial roles in signal transduction and various developmental and stress response processes.^69,71,73^ Notably, Lehti-Shiu et al.^17^ did not detect the AGC_PKA-PKG, BUB, PEK_PEK, and TKL-PL-8 families in the cucumber species; however, our study identified their presence. Discrepancies in analytical methodologies and the absence of genome collinearity among the studied cucumber lineages may account for these differences.

According to Lehti-Shiu et al.^17^, the 20 PK groups detected in our study could be subdivided into 123 families, with 34 of them composed of only one PK gene. Notably, the RLK-Pelle_DLSV family was the most numerous, consisting of 68 proteins, a result compatible with that obtained for the cucumber species by Lehti-Shiu et al.^17^. Similar results have also been found in other species, such as Gossypium,^78^ rice,^79^ Arabidopsis^67,68^ and corn.^70^ The expansion and establishment of this family probably occurred before the divergence of Viridiplantae species,^17^ as no evidence that the cucumber species has undergone whole genome duplication.^16^ However, after a new assembly of the cucumber genome, Li et al.^42^ reported 239 tandemly duplicated genes, and Yu et al.,^80^ reporting on 15 pairs of tandem duplication genes and 31 pairs of segmental duplication in the cucumber genome.

In addition to the RLK-Pelle_DLSV family, another 23 subfamilies, comprising 202 proteins with LRR domains, also stood out in the RLKs group due to their wide occurrence. These findings align with those presented by Yu et al.^79^, who identified 189 genes distributed across 22 families RLK-LRR in the Chinese Long cucumber genome v3. The substantial presence of genes within this family underscores their significance in stress response mechanisms, as highlighted in studies by Dambroz et al.,^81^ Fischer et al.,^82^ Magalhães et al.,^83^ and Sakamoto et al.^84^

### 4.2. Characterization of the cucumber PK sequences

Addressing diverse characteristics of genes and the proteins they encode provided a broad insight into the structural and functional diversity of cucumber PKs. Initially, we sought to explore the structural attributes of the 835 PK genes, such as determining the number of introns. The presence of introns may be intrinsically associated with regulating gene expression, as in processes such as alternative splicing. Notably, intronic genes, which encompass specific intron fragments undergoing expression and factors such as intron length and intron phase, assume crucial roles in the regulation of gene expression.^85^ Researches report the presence of coding genes inside intronic regions.^86,87^ These findings underscore the structural diversity inherent in cucumber’s PK genes and shed light on the potential intricacies of their regulatory landscapes.^88^ The presence of introns in a gene can influence its susceptibility to silencing by microRNAs (miRNAs). The miRNAs can bind to mRNA sequences that include introns silencing them, and the number of introns can affect the efficiency of miRNA-mediated silencing.^89,90^ Introns play a role in the regulatory process and the number of introns can bring crucial inferences. The presence of introns in miRNA target genes can impact mRNA maturation and translation efficiency, thereby influencing overall gene expression dynamics.^90^

The uneven distribution of PK genes on the seven chromosomes of the cucumber genome represents a non-random pattern, which may be a result of the evolutionary process undergone by the species.^63^ However, the distribution of RLK genes in clusters was observed, supporting the hypothesis that this family originated by tandem duplication and later subfunctionalization,^91^ although no duplicatin was evidenced in cucumber species.^16^ On the other hand, the results of Li et al.,^42^ and Yu et al.^80^ contradict previous studies, giving evidence of the cucumber genome duplication. These conflicting findings highlight the complexity of the cucumber genome’s evolutionary history, suggesting that further research is needed to clarify the mechanisms underlying gene distribution and duplication in this species.

Considerable variation was observed in all the characteristics of PKs examined, indicating specialized functions in their respective microenvironments, similar to patterns observed by Dambroz et al.^81^ Most of the functions performed by PKs are associated with the plasma membrane and nucleus, comprising about 77% of the PK genes in cucumber. It has been observed that the kinase domains of nuclear and membrane proteins have been associated with pathogen identification and plant defense responses. The predominant functions of PKs associated with the plasma membrane and the nucleus include the perception and response to environmental stimuli, such as abiotic stress and resistance to pathogens, as well as the regulation of subsequent gene expression.^92,93^ This dual functionality, spanning both nuclear and membrane realms, underscores the versatility of PKs in performing essential cellular processes.^94^ At the plasma membrane, PKs are instrumental in perceiving external signals, transducing them into intracellular responses, and modulating various physiological pathways crucial for plant growth and adaptation.^95^ Simultaneously, within the nucleus, PKs contribute significantly to the intricate machinery of gene regulation, influencing transcriptional processes that ultimately shape the plant’s responses to environmental challenges.^96^ A pattern of specialized functions localized predominantly in the plasma membrane and nucleus emerges, aligning with observations in cucumber, common beans,^68^ *Hevea brasiliensis* and *Manihot esculenta*.^97^ This consistency across diverse plant species suggests a fundamental role for PKs in essential cellular processes and stress responses. However, the observed diversity in subcellular locations among PKs also hints at their involvement in specific signaling pathways, highlighting the versatile and intricate nature of their functions. Similar results were also observed in common beans,^69^ *H. brasiliensis*, and *M. esculenta.*^91^ However, other genes can be found in various subcellular locations, suggesting their involvement in specific cellular processes, and signaling pathways in various cellular biological processes together with the stress responses.

In principle, PKs are responsible for phosphorylation processes and binding to specific targets in the cell related to activation processes. Therefore, they can be listed as interesting targets for genetic breeding, considering the hypothesis of improvement of central metabolic processes, such as increased efficiency of enzymatic conversion or modifications at the binding site that recognizes pathogen’s effectors, preventing its infection. As an example, regulation of gene expression through transcription factors led to overexpression of mutant rice genotypes, increasing nitrogen use efficiency and productivity.^98^ However, studies detailing these processes involving phosphorylation remain scarce.

### 4.3. Kinase gene expression patterns

The study of differential gene expression has been one of the most robust and sophisticated methodologies for understanding the central nuances of pathogen-plant interaction (PPI). This type of interaction involves several processes including the perception of extracellular signals, the phosphorylation of specific proteins, and the expression of various defense-related genes and tissue remodeling. In this study, we investigated the expression profiles of the 22,626 genes encoded in Gy14 genome in response to biotic stress caused by *P. xanthii* using transcriptome data,^24^ enriching our analysis for 835 genes encoding protein kinases.

Our analysis identified PK DEGs in response to *P. xanthii* infection (PM), showing similar expression patterns but at different response times. Some DEGs act at the cell nucleus level (∼25%) in processes related to protein phosphorylation and mitotic cell cycle regulation; while most of the DEGs (∼55%) function in the plasma membrane, with general functions related to phosphorylation and participation in some more specific processes such as Peptidyl-Tyrosine Phosphorylation, MAPK Cascade, Hormone-Mediated Signaling Pathway, Phragmoplast Assembly. Some proteins in smaller numbers were also located in chloroplasts, endomembrane system, extracellular space, mitochondrion, and mitochondrial membrane with functions related to Polysaccharide Binding, Nucleotide Binding, Transmembrane Receptor Protein Tyrosine Kinase Activity, Polysaccharide Binding; and metabolic processes, such as Defense Response to Bacterium, and Peptidyl-Tyrosine Phosphorylation. These findings suggest that PKs play important roles in the defense response against *P. xanthii*, with similar expression patterns observed between resistant and susceptible genotypes and major expression differences related to response time to infection,^99^ because in general, susceptible genotype shows an immediate response, while the resistant genotype show a later response after infection. The shallow response time may be related to the time of cell maturation, time of development of infection-induced apoptosis, rates of infection spread, and recovery of infected cells. In addition, these factors may express themselves with different intensity among different genotypes, suggesting that uninfected cells do not have sufficient acquired immunity to recover the host from infection.^100^ In this way, resistant genotypes have a more effective immune response responding more quickly against infection, due to several factors including localized immune efficacy.^101,102^ The difference in stress response time also involves the perception of changes in the environment by plant tissues and directly interferes with the cell cycle. This perception involves the activation of CDKs, which are the central regulators of this cycle. Cyclin-dependent kinases (CDKs) play important roles as it relates to cell division and response to many intracellular and extracellular signals.^103^ This process results in the activation of a signaling cascade that prolongs the S phase with delayed entry into mitosis.^104^ This reveals the importance of the PKs of the CMGC families to cucumber resistance to *P. xanthii*.

While the predominance of RLKs among DEGs associated with resistance reinforces the importance of these receptors in perceiving pathogen signals and triggering defense responses, as observed in other plant species, the presence of MAPKs and CAMKs suggests the activation of kinase-mediated signaling pathways, known to orchestrate responses such as ROS production and the hypersensitive response. Their subcellular localization predominantly in the plasma membrane and nucleus reinforces this role in perceiving signals and regulating subsequent gene expression.

### 4.4. Kinase duplication and synteny analyses

Our detailed analysis of the duplication events revealed significant insights into the cucumber kinome. The estimates performed for 833 kinases identified the predominance of doubling events, highlighting notable contributions of total genomic duplications (WGD) and tandem duplications. The predominance of WGD events, which account for approximately 69.4% of kinases in cucumber, suggests a key role of these duplications in the evolution of the kinome. These events can be associated with specific spikes in synonymous substitution rates (Ks), indicating significant duplications around 50.41 and 422.81 million years ago (MYA). Notably, this last interval coincides with polyploidization events in the eudicot lineage, pointing to a possible influence of these events on the diversification of these kinases.

Analysis of the non-synonymous substitution rate (Ka) revealed that most kinases in cucumber are under purifying selection, indicating an evolutionary pressure to preserve the structure and stabilize the function of these proteins. This pattern is consistent with other studies on different gene families in common beans,^69^ suggesting a general trend in functional conservation across the genome. The identification of 237 kinases with tandem duplications is especially interesting. Among these, notable families, such as RLK-Pelle_DLSV, exhibited a significant number of protein duplications. These tandem duplications are known to be associated with stress responses, indicating a possible adaptive expansion of these subfamilies to address environmental challenges.^105,106^

When comparing our results with studies in other plant species, we observed similar patterns regarding the presence of duplications under purifying selection. However, we highlight the uniqueness of our findings, especially in terms of high rates of Ks and distant dates of doubling for the kinases. These traits may be specific to the evolution of kinases in this species and may influence functional diversity and long-term adaptation.

We underscore the pivotal role of Protein Kinases (PKs) in cucumber immunity and disease resistance. Our comprehensive analysis delineates the characteristics of cucumber’s 835 PKs, categorized into 123 families and 20 groups. Notably, only 312 PKs demonstrated significant relevance in response to biotic stress, further classified into 10 groups: AGC, BUB, CAMK, CMGC, RLK, STE, TKL, ULK, WEE, and WNK. This investigation establishes an important groundwork for future exploration into the specific functional roles of these PKs, paving the way for the development of targeted strategies to enhance disease resistance in cucumbers and other Cucurbitaceae species. The outcomes of this research contribute significantly to the progression of agricultural practices, offering valuable insights for bolstering crop resilience against diverse disease challenges.

## Abbreviations

aa: (amino acid)
AGC: (cAMP-dependent protein kinases cGMP-dependent protein kinases various types of protein kinase C protein kinase B 3-phosphoinositide-dependent protein kinase-1 and the ribosomal protein S6 kinases)
ALS: (Alternaria Leaf Spot)
Aur: (aurora kinase)
BUB: (budding uninhibited by benzimidazoles)
C: (cellular component)
CAMK: (calcium/calmodulin-dependent protein kinase)
CDK: (Cyclin-dependent kinase)
CDPK: (Calcium-Dependent Protein Kinase)
CK1: (casein kinase 1)
CMGC: (cyclin-dependent kinase mitogen-activated protein kinase glycogen synthase kinase and cyclin-dependent-like kinase families)
CPU: (central processor unit)
CRK: (CDPK-Related Protein Kinase)
Csa: (Cucumis sativus)
CuGenDbv2: (Cucurbit Genomics Database v2)
DEG: (Differentially Expressed Gene)
F: (molecular function)
FDR: (false discovery rate)
Gene Ontology: (GO)
GRAVY: (Grand Average of Hydropathy)
HMM: (Hidden Markov Model)
iP: (Isoelectric Point)
IRE1: (inositol-requiring enzyme 1)
Ka: (Non-synonymous substitution rates)
Ks: (Synonymous substitution rates)
LecRLK: (Lectin Receptor-Like Kinase)
log2fc: (log2 fold change)
LRR: (Leucine Rich Repeat)
MAPK: (Mitogen-Activated Protein Kinase)
MYA: (million years ago)
NAK: (NF-κB-activating kinase)
NEK: (never in mitosis gene-A)
P: (biological process)
PEK: (pancreatic eukaryotic initiation factor 2 α-subunit kinase)
PK: (Protein Kinase)
Pkinase: (Protein Serine/Threonine Kinase)
Pkinase_Tyr: (Protein Tyrosine Kinase)
Plant specific: (Group-Pl)
PM: (Powdery Mildew)
PPI: (pathogen-plant interaction)
RKN: (Root-Knot Nematode)
RLK: (receptor-like kinase)
RNA-Seq: (RNA sequencing)
SCY: (Saccharomyces cerevisiae [yeast] kinase)
STE: (serine/threonine kinase)
TKL: (tyrosine kinase-like kinase)
TLK: (tousled-like kinase)
TM: (transmembrane domain)
TTK: (threonine/tyrosine kinase)
ULK: (unc-51-like kinase)
WAG: (Whelan Goldman)
WEE: (wee1 wee2 and myt1 kinases)
WGD: (Whole genome duplication)
WNK: (with no lysine-K)

## Data availability

Data in pre-publication were analyzed in this study relative to the *Cucumis sativus* Gy14 genome with permission for publication from Dr. Yiqun Weng (USDA-ARS Vegetable Crops Research Unit). The data from the *Cucumis sativus* Gy14 genome, and public data of RNA-seq project PRJNA321023 (PM) are available in the Cucurbit Genomics Databsase v2 (CuGenDBv2) (http://cucurbitgenomics.org/v2/).

## Conflict of Interest

The authors assert that they do not possess any identifiable financial conflicts of interest or personal associations that might have seemed to impact the findings presented in this manuscript.

## Acknowledgements

This work was supported by grants from the Coordenação de Aperfeiçoamento de Pessoal de Nível Superior (CAPES) – Finance Code 001, and Fundação de Amparo à Pesquisa do Estado de Minas Gerais (FAPEMIG). FC received a DSc fellowship from CAPES (88887.480460/2020-00), and WP received support from Universal Demand, Process n. APQ-00511-23. Our sincere gratitude to Dr. Yiqun Weng for gently providing the data from the Gy14 genome in pre-publication for this research.

## Supporting Information

Supplementary Table S1. Kinase domain annotation of the 835 cucumber protein kinases (best hit).

Supplementary Table S2. Domain annotation of the 835 cucumber protein kinases

Supplementary Table S3. Predicted domain combinations for each of the 835 cucumber protein kinases

Supplementary Table S4. Family kinase classification

Supplementary Table S5. Kinase subfamily and group quantification

Supplementary Table S6. Gene structure analysis.

Supplementary Table S7. Kinase characterization

Supplementary Table S8. Gene Ontology (GO) of protein kinases of cucumber

Supplementary Table S9. Differentially expressed genes encoding protein kinases and their classification as resistance, susceptibility or unrelated genes to Powdery Mildew.

Supplementary Table S10. Duplication analysis

Supplementary Table S11. Synteny analysis

## Notes

### Competing Interest Statement

The authors have declared no competing interest.

### Summary of Updates

In this review, we kept the kinome analysis but focused on the powdery mildew stress, removing information from the Alternaria leaf spot and Root-knot nematode.

