## Supplementary figures and images for "The *Cucumis sativus* kinome: Identification, annotation, and expression patterns in response to powdery mildew infection"

### Supplementary figure S1

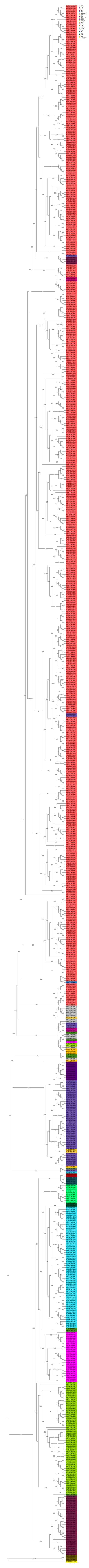
